# Transcriptional output, cell types densities and normalization in spatial transcriptomics

**DOI:** 10.1101/503870

**Authors:** Manuel Saiselet, Joël Rodrigues-Vitória, Adrien Tourneur, Ligia Craciun, Alex Spinette, Denis Larsimont, Guy Andry, Joakim Lundeberg, Carine Maenhaut, Vincent Detours

## Abstract

Spatial transcriptomics measures mRNA at hundreds of 100 micrometer-diameter spots evenly spread across 6.5×6.9 mm2 histological slices. Gene expression within each spot is commonly normalized by total read counts. However we show that the transcriptional output of individual spots reflects the number of cells they contain, hence total read counts per spot reflect relevant biology. Although per-spot read-count normalization reveals important enrichment trends, it may heavily distort cell-type-related absolute local expression and conceal important biological information.

## Dear editor

Spatial transcriptomics (ST) bridges untargeted genome-wide mRNA profiling with tissue morphology by making it possible to perform RNA-seq at hundreds of precisely located spots on the surface of an histological slice (Ståhl et al., 2016). Since mRNA diffusion is minimal during tissues permeabilization and mRNA capture, the transcriptome of each spot is thought to aggregate the transcriptomes of the cells it contains. The number of cells within a spot and their transcriptional output depend on their type, state and on overall local morphology. ST data share many limitations of single cell RNA-seq, including low coverage and high dropout rate. So far, ST studies have relied on preprocessing pipelines inspired by single cell RNA-seq studies (Ståhl et al., 2016; Asp et al., 2017; Giacomello et al., 2017; Berglund et al., 2018; Lundmark et al., 2018; Salmen et al., 2018; Thrane et al., 2018). These include normalization of gene-wise read counts in a cell/spot by the total number of reads collected from that cell/spot. But the number of reads obtained from a spot could reflect its cellular content or technical variation in RNA capture and amplification. Thus, whether read count normalization is warranted in the context of ST remains an open question. We addressed it by quantifying the cellular content of individual spots from image analysis and by comparing it with read counts.

A BRAF V600E-mutated papillary thyroid cancer (PTC) was profiled with ST (Supplementary Material and Methods). Pathology review of the H&E image revealed five major types of morphologies (Fig. 1A). This qualitative approach was complemented by whole-slide machine learning-based localization of nuclei, and their classification within three categories (Fig. 1B): epithelial cells, fibroblasts and ‘other cells’, which mostly contains immune cells. Among 86,111 detected nuclei (Fig. 1C), 31% were located within ST spots. The mean number of cells per spots varied from 0 to 197 (median 67).

**Figure 1.**
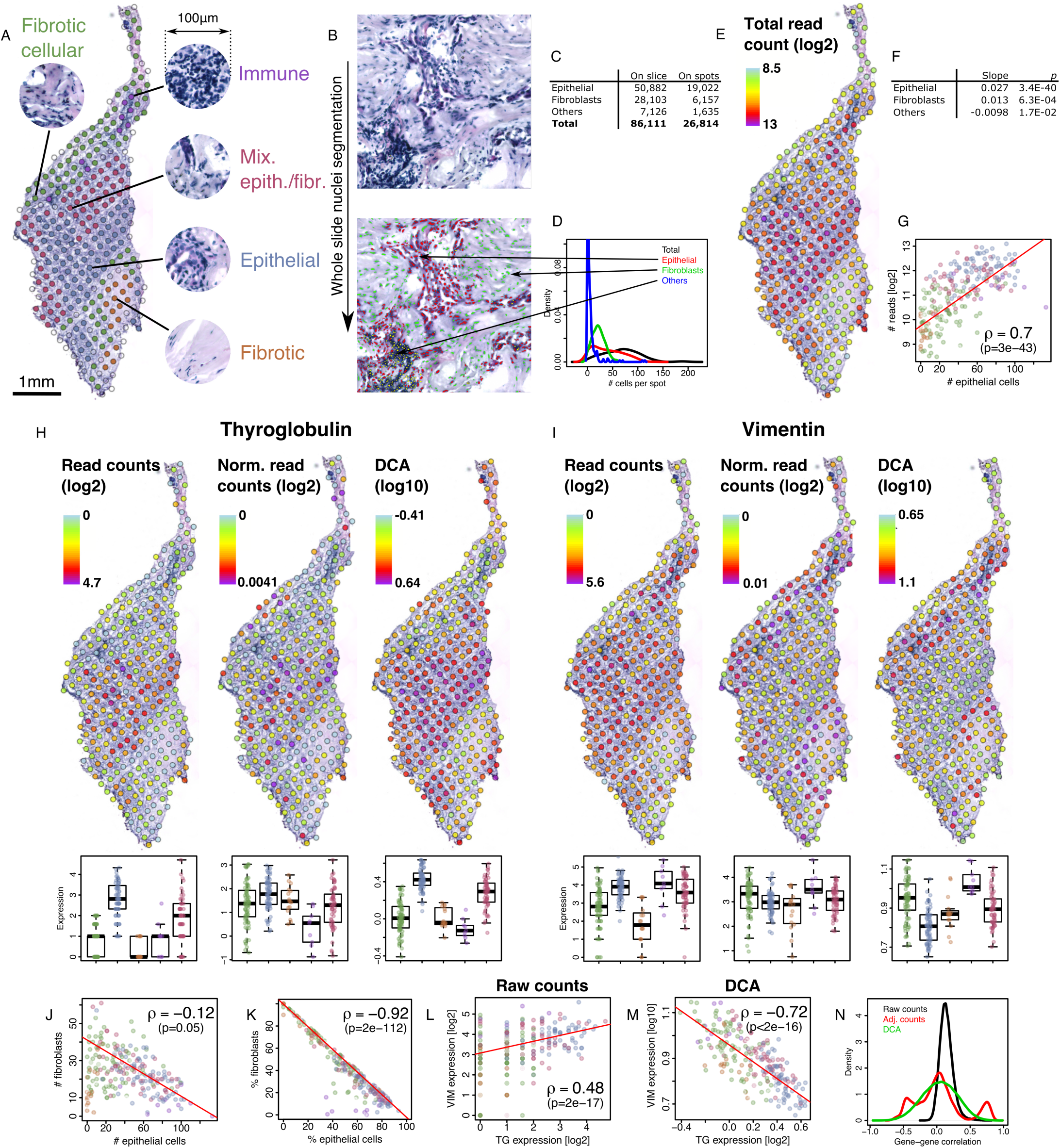
Variation of total read counts is related to morphology and number of cells of different types. **A**, Five types of morphological regions were determined from pathology. The transcriptome was determined for each spot with ST. **B**, The nuclei on the H&E image were segmented and classified with a machine learning-based algorithm (see Supplementary Material and Methods). **C**, Nuclei counts. **D**, Distributions of the number of cells per ST spot. **E**, Total read count per spot. **F**, Multivariate analysis of association between cell numbers and the log_2_ of total read count per spot. **G**, Each point represents a ST spot with same color code as in panel A, ρ denotes the Spearman’s correlation. **H**, Expression of thyroglobulin without normalization (left), with adjustment for total read counts (center), and with a neural autoencoder-based normalization (DCA, right). Boxplots represents the expression of spots in the regions shown in panel A (same color code). **I**, Same as panel **H**, except that vimentin expression is shown. **J**, Points represent spots (same color code as in panel A) with the absolute number of epithelial cells (*x*-axis) and the fibroblasts (*y*-axis). **K**, Same as **J**, except that cell types proportions are depicted. The large negative correlation stems from the low number of ‘other’ cells (panel C). **L** and **M**, Expression of TG and VIM are compared using raw counts or auto-encoder-based normalization. **N**, Distribution of gene vs. gene correlation across spots for raw counts and normalized data.

Spot-wise read coverage varied from 356 to 8,749 across the slide (Fig. 1E), a 25-fold variation. It was associated with the number of cells of all types in a multivariate analysis (Fig. 1G), particularly epithelial cells (Fig. 1F). As expected, total read count per spots was highest in dense epithelial areas, and lowest in low cell density fibrotic zones (Fig. 1E,G). We concluded that total read counts per spot reflects relevant quantitative and qualitative features of tissue morphology.

To assess the effect of normalization we compared raw read counts with raw counts normalized for total counts and with scale-normalized expression estimates generated by DCA (Eraslan et al., 2019) a recent neural autoencoder-based algorithm developed in the context of single cell RNA-seq. Figs. 1H and 1I show expression of thyroglobulin (TG, a thyroid differentiation marker) and of vimentin (VIM, a mesenchymal intermediate filament), respectively. Raw counts and normalized expression of TG all closely followed epithelial density and total counts (compare Fig. 1H to Figs. 1A and 1E, see also Supplementary Fig. 1). VIM raw counts were substantial in the epithelial areas, but also in the cellular fibrosis and immune foci. Normalization, however, revealed a dramatically different picture, particularly for DCA: while remaining high in cellular fibrosis and immune foci, VIM expression was lower in epithelium (Fig. 1I, Supplementary Fig. 1).

Thus, normalization affected the spatial expression pattern of VIM, but not TG. The absolute numbers of epithelial cells and fibroblasts per spot were weakly associated (Fig. 1J), while their relative proportions, *i.e.* their number divided by the total number of cells within a spot, were massively anti-correlated (Fig. 1K). The raw counts of TG and VIM are positively correlated (Fig. 1L), while their normalized values are negatively correlated (Fig. 1M). The positive correlation between TG and VIM raw counts (Fig. 1L) suggests that the tumoral epithelium expresses VIM and could undergo an epithelial-mesenchymal transition. This transition has been reported in BRAF V600E-mutated tumors (Knauf et al., 2011) such as this one. VIM is also expressed in primary cultures of normal thyrocytes treated with Epidermal Growth Factor, which inhibits differentiation, but not of thyrocytes treated with Thyroid Stimulating Hormone, which promotes thyroid differentiation—while both are mitogenic (Coclet et al., 1991). Alternatively, VIM expression by fibroblast could be promoted by nearby epithelial cells. Overall, raw counts seemed more related to the number of cells of a given cell type, while normalized expression captured cell types’ relative proportions.

To rule out possible artifacts related to TG and VIM in a particular tissue slice, we reproduced the above analysis 1-to the thyroid stimulating hormone receptor (TSHR) and collagen III α1 (COL3A1) in another slice of the same thyroid cancer (Supplementary Fig. 2); 2-to the estrogen receptor (ESR1) and VIM in a publicly available breast cancer slice profiled on the recent Visium platform (10X Genomics, Pleasanton, USA; Supplementary Fig. 3). Taken together, these controls establish the generality of the effect of normalization on the relation between epithelial differentiation and mesenchymal markers, and their relevance to Visium slides, which have a 4-fold higher resolution than the first generation ST slides of Fig. 1.

To gain insights on the global effect of normalization, we calculated the distribution of correlations between genes across spots for all three expression metrics (Fig. 1N). Raw counts were positively correlated for most pairs of genes. By contrast, normalized expressions correlations were centered on 0. This implies that genes tend to show a similar expression pattern that reflects total transcriptional output when raw counts are considered, while normalized expression highlights contrasts between genes.

We showed that the variation of total read counts is largely determined by local cell density in ST data. Thus, total counts per spot are biologically informative and do not necessarily need to be normalized out. Some single cell analysis pipelines tie normalization and denoising, yet these are technically independent operations. For example, DCA estimates read counts scale factors, but users are free to use its scale-adjusted or un-adjusted outputs (Eraslan et al., 2019). Our study shows that both options are valid, but address different purposes.

Raw read counts inform about the absolute density of cell types, while normalized expression informs about their relative proportions. It is remarkable that normalized expression better detects specific morphologies such as pure epithelium and cellular fibrosis (see boxplots Figs. 1H and 1I, compare the DCA panels of Figs. 1H and 1I with the pathology annotation of Fig.1A), while raw counts do reflect actual cell-type local densities and may highlight atypical expression patterns, as exemplified here for VIM in regions of high epithelial density.

The resolution of commercially available spatial transcriptomics will eventually reach sub-cellular resolution (Vickovic et al., 2019). Given that some cells, e.g. cancer cells, produce more RNA that others (Lovén et al., 2012), it begs the question of to what extent our argument also applies at single cell level. Cell level phenotypic information measured independently of transcription must be available together with matched cell transcriptomes in order to unambiguously address this question.

## Author contributions

M.S. performed experiments with support of J.L.. J.R.V. and V.D. performed computational analysis, L.C., A.S. and D.L. handled sample banking and pathology review, G.A. resected the tumor. C.M. and V.D. supervised the research and wrote the manuscript.

## Acknowledgments

We thank Annelie Mollbrink and Jose Navarro for help with the ST protocol and bioinformatics.

## Supplementary data

Supplementary Material and Methods are available online on *JMBC* web side. Code and data are available at https://github.com/vdet/st-normalization.

## Supplementary Material and Methods

### Code and data availability

Code and data are available on GitHub: https://github.com/vdet/st-normalization

### Sample

The sample is a pT1a stage, BRAFV600E-mutated, papillary thyroid cancer (PTC) removed from 40 years old female at the Institut Jules Bordet. This patient was diagnosed with hypothyroidy at age 15 in the context of a simple goiter. At 22, a small cystic and hypoechogen nodule was detected in the right para-isthmic region of her thyroid. At 35, an hypoechogen nodule <1cm is detected in the left lobe. It remained stable over the following year at 7×7×8mm^3^. At 40, a fine needle biopsy revealed a PTC, which was removed. No peripheral adenopathy was detected by echography. This study was approved by the ethics committee of the Jules Bordet Institute (protocol #1978).

The resected tissue was immediately dissected, placed on ice, embedded in preservative solution (OCT), frozen and stored at −80°C. The specimen was sliced at 10µm thickness with a cryostat. Six consecutive slices were processed on a library preparation microarray ST slide.

Total RNA was extracted from tissue slices using Qiazol, followed by purification on miRNeasy columns (Qiagen) according to the manufacturer’s recommendations. A RNA integrity quality score of 9.0 was defined using an Experion (Bio-rad) according to manufacturer’s recommendations.

BRAF mutational status was established according to a previously described methods.^1^ BRAF V600E positive and negative samples previously obtained in the lab were used as controls.

### Spatial transcriptomics

ST profiling was performed as described by Salmén *et al.*^2^ For our sample, hematoxylin incubation was performed during 7 minutes followed by eosin incubation during 60 seconds. Pepsin/HCl permeabilization was performed during 10 minutes and a 3X Beta-Mercaptoethanol protocol was applied during one hour in order to remove residual tissue.

### Nuclei segmentation and classification

The nuclei segmentation model was established on the basis of 9 images of 1,024×1,024 pixels cropped from the original 30,131×27,755 H&E image. First, we interactively trained a pixel learning algorithm, iLastik v1.3.0 pixel classification module,^2^ with the purpose of labeling pixels in four classes: empty space, fibrosis, epithelial cytoplasm and nuclei. Many examples of cytoplasmic pixels between nuclei were provided in order to resolve areas with densely packed nuclei. Second, starting from the nuclei masks, we trained the iLastik object classification module to recognize epithelial nuclei, fibroblasts and cells that are neither epithelial, nor fibroblasts. The two classifiers were then applied to the entire image. The H&E image and resulting masks are available from the companion GitHub page. Nuclei positions were computed with the ‘`Analyze Particules`’ function of Fiji v2.0.0,^3^ requesting at least 4 pixels per nuclei.

ST spot coordinates were corrected with `st_spot_detector`^4^ as described in ref. 5. The resulting corrected coordinates were used for subsequent graphical visualization of the data as well as calculation of spot-wise nuclei counts. A total of 1,575 spots were positioned on-tissue across the 6 serial slices.

### Computing raw read counts from sequence data

The FASTQ files obtained by Illumina sequencing were processed using `st_pipeline` v1.6.0 as described in ref. 5, with default parameters and using the 1000L8 spatial barcode reference file. Alignments rested on STAR v2.5.4,^6^ the human reference genome Hg38 and the Genecode v27 gene model. `st_pipeline` outputs for each tissue slice the matrix of UMI-collapsed raw counts, per gene, per spot.

### Gene filtering and read counts normalization

Three filters were applied to discard genes with low counts. First, we eliminated any gene with higher read counts in spots outside the tissue area than in spots covering the tissue, on a per slice basis. Second, we removed any gene not expressed on all six consecutive tissue slices we profiled with ST. Third, we ranked genes by increasing total count across the 6 slices and kept for further analysis the most expressed genes that, taken together, represented 80% of the total read mass. Formally, let *c*_*i, j, k*_ be the number of read aligned on gene *i* for spot *j* of slide *k*. We ranked genes by decreasing Σ_*j, k*_ *c*_*i, j, k*_, and kept for subsequent analysis genes with the rank *r<R*, with *R* set such that Σ_*i* ≤*R, j, k*_ *c*_*i, j, k*_ /Σ_*i, j, k*_ *c*_*i, j, k*_ ≤ 0.8. This filter has the potential to remove genes with high expression in a small number of spot. This, however, was not a major issue as only two genes with a maximal read count greater than 10 were filtered out. In the end, 3,535 genes were used in subsequent analyses.

To normalize for total count we divided count values for each spot by the total number of reads for that spot. These calculations were carried out with R v3.4.4 on a per slice basis.

The Deep Count Autoencoder^7^ code was downloaded from https://github.com/theislab/dca on 04/26/2018 and run with default parameters on the raw count matrix of 1,575 spots and 3,535 genes.

### Processing of Visium breast cancer data

Data were downloaded from https://www.10xgenomics.com/resources/datasets/, we used sector 2 for visualization. The DCA algorithm was run with default parameters, using concatenated raw counts from sector 1 and sector 2. Data were read into R and Displayed with the Seurat R package v3.1.2.

## Supplementary Figures

**Supplementary figure 1.**
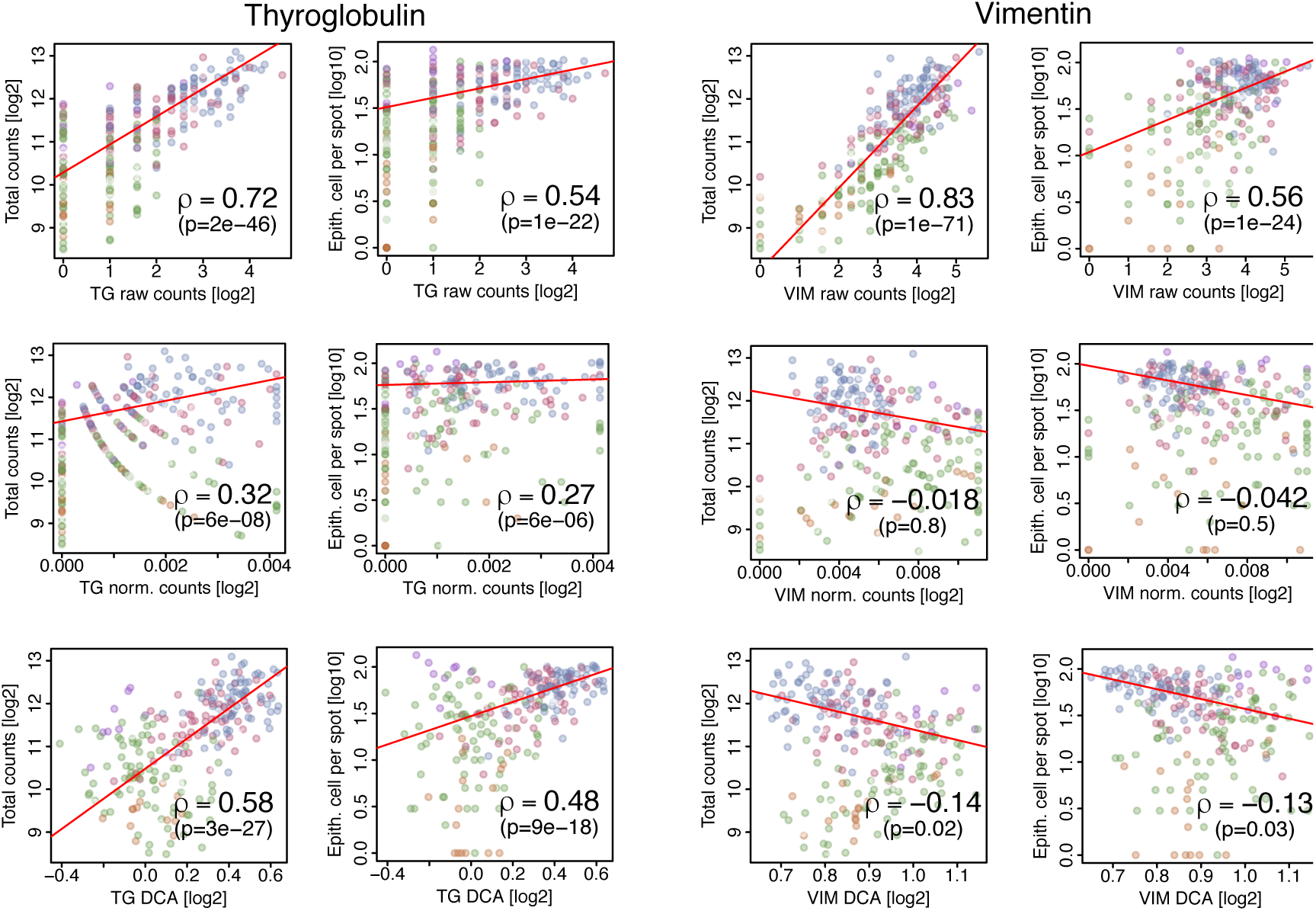
Effect of normalization on correlations of TG and VIM with total counts and epithelial cell density.

**Supplementary figure 2.**
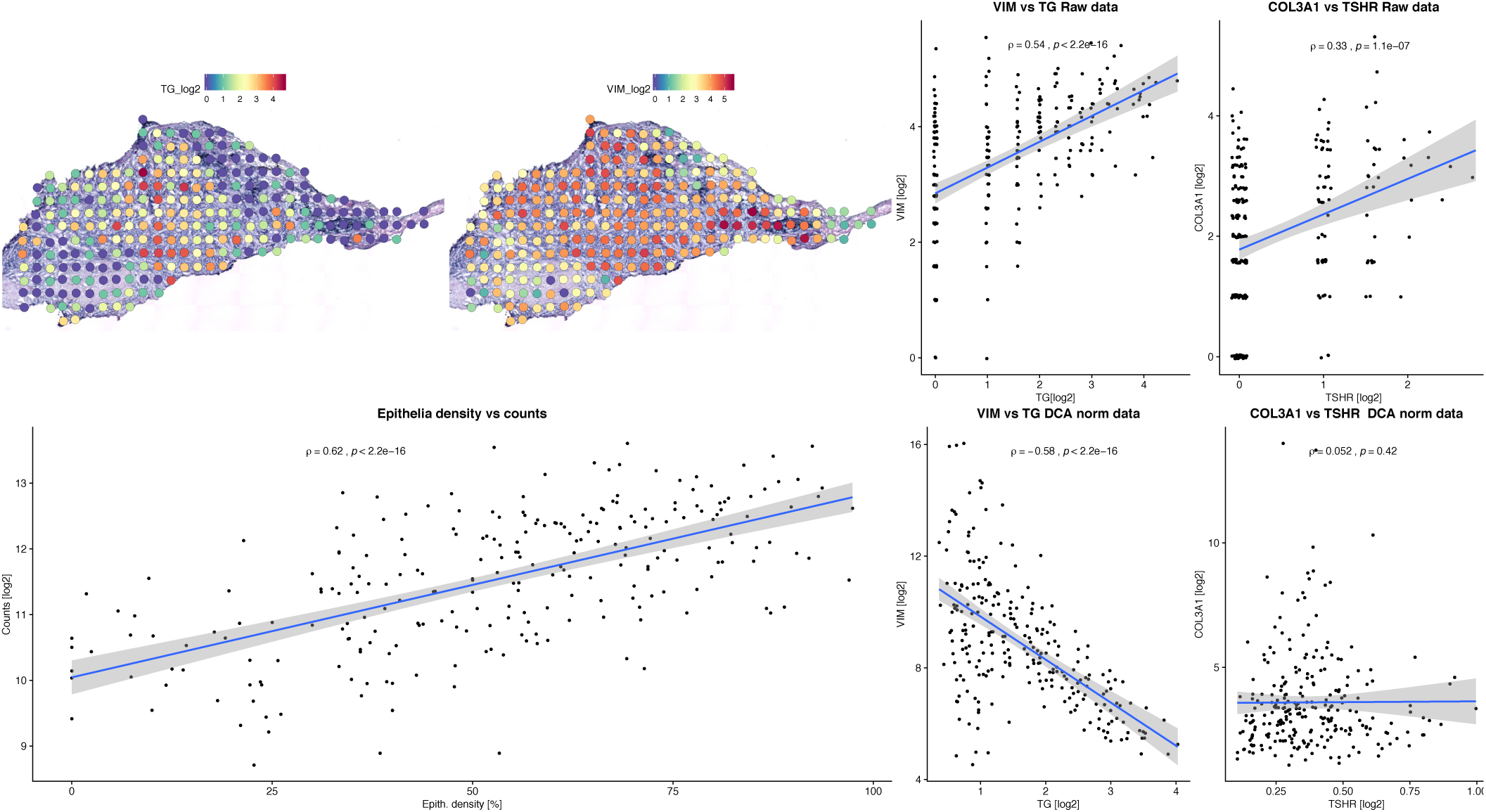
We reproduced the analysis of Fig. 1 on another tissue slice cut from the same tumor. The heatmaps show the raw counts for TG and VIM. The left-most scatter plot shows that total read counts is correlated with the density of epithelial cells. The rightmost panels show the correlation between TG and VIM or TSHR and COL3A1 computed from raw counts of DCA-normalized data. TSHR is another major thyroid follicular cell marker. COL3A1 is an extracellular matrix component that is believed to be expressed by fibroblasts.

**Supplementary figure 3.**
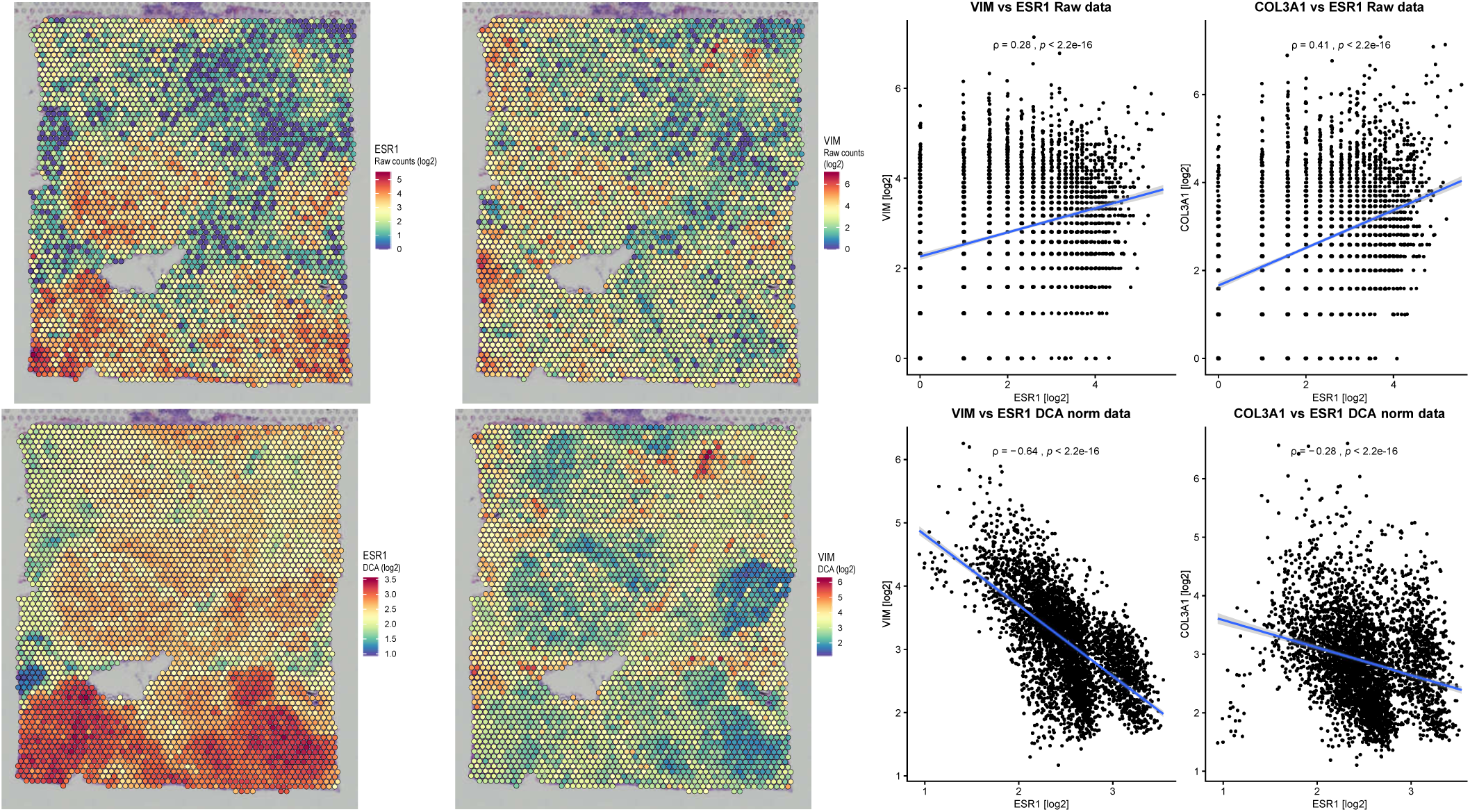
The top heatmaps represent the raw counts for ESR1 and VIM across a breast cancer slice profiled with the Visium Platform (10X Genomics). The bottom heatmaps show the estimated expression of ESR1 and VIM after DCA denoising and normalization. Note that the variation of total counts across this slice is >20-folds (not shown). Normalization reverses the correlations between ESR1 and VIM, and between ESR1 and COL3A1 (scatter plots on the right). The estrogen receptor 1 (ESR1) is a differentiation marker of the breast epithelium, i.e. its role in this analysis is analogous to the role of TG in Fig. 1 and TSHR in Supplementary Fig. 2.

